# Integrative analysis of TCGA transcriptomic states and DepMap dependencies prioritizes candidate vulnerabilities in immune-cold microsatellite-stable colorectal cancer

**DOI:** 10.64898/2026.07.04.736484

**Authors:** Amol Tandon, Deepthi Nagalla

**Affiliations:** Department of Biochemistry and Molecular Genetics, University of Alabama at Birmingham, Birmingham, Alabama, United States of America; German Cancer Research Center (DKFZ), 69120, Heidelberg, Germany; SOSS, Indira Gandhi National Open University (IGNOU), India

**Keywords:** colorectal cancer, MSS/MSI-L, immune-cold tumour microenvironment, TCGA, DepMap, cancer dependency, ERBB2, HER2, immune resistance

## Abstract

Microsatellite-stable/microsatellite instability-low colorectal cancer (MSS/MSI-L CRC) is generally resistant to immune checkpoint blockade, but the biological states underlying this resistance are heterogeneous. We integrated TCGA COAD/READ patient transcriptomic profiles, MSIsensor-based MSS/MSI-L classification, curated immune and stromal module scoring, focused differential expression and DepMap CRISPR dependency data to prioritize candidate vulnerabilities in immune-cold MSS CRC. Among 494 MSS/MSI-L tumours, 218 were classified as MSS intermediate, 102 as MSS immune-cold, 91 as MSS hot/inflamed and 83 as MSS barrier-high. MSS immune-cold tumours showed lower cytotoxic, IFNγ-chemokine and antigen-presentation programmes than MSS hot/inflamed tumours, including reduced NKG7, CD8A, CXCL9, CXCL10 and LAG3 expression. MSS barrier-high tumours showed enrichment of stromal and extracellular-matrix programmes, including COL1A1, COL1A2 and COL3A1. Integration with DepMap CRISPR gene-effect data from 1208 cancer models, including 63 colorectal cancer models, separated tumour-cell-intrinsic dependencies from patient-derived microenvironmental signatures. Candidate target classes included ERBB2, VEGFA, PIK3CB, ATR/WEE1/CHEK1, HDAC1/HDAC3/BRD4 and BCL2L1/MCL1, while collagen genes were interpreted as stromal-barrier markers rather than tumour-cell dependencies. ERBB2 expression was higher in MSS immune-cold than MSS hot/inflamed tumours and further elevated in MSS barrier-high tumours, supporting ERBB2 as a candidate subset-associated signal that requires orthogonal HER2 validation. These findings support a stratified therapeutic framework for immune-cold, barrier-high and intermediate MSS CRC.

## Introduction

Colorectal cancer (CRC) remains a major global health burden and is among the leading causes of cancer-related mortality worldwide. Although molecular stratification has improved the management of selected CRC subgroups, advanced disease remains difficult to treat, particularly after progression on standard chemotherapy and targeted therapy. Immune checkpoint blockade has transformed the treatment of several solid tumours and has produced durable responses in mismatch repair-deficient or microsatellite instability-high (dMMR/MSI-H) CRC. However, the majority of CRCs are mismatch repair-proficient and microsatellite-stable or MSI-low (pMMR/MSS/MSI-L), and these tumours usually derive limited benefit from immune checkpoint inhibitors when used as monotherapy [1–4].

The clinical contrast between MSI-H and MSS CRC has become a central problem in gastrointestinal immuno-oncology. MSI-H/dMMR tumours typically carry higher tumour mutational burden, increased neoantigen load, stronger immune infiltration and more prominent interferon-associated inflammatory programmes. These features provide a biological basis for responsiveness to PD-1-based therapy. In contrast, MSS CRC frequently exhibits reduced T-cell infiltration, weak cytotoxic immune activity, impaired inflammatory chemokine signalling, stromal or myeloid-mediated suppression, tumour-intrinsic immune-evasion pathways and lower baseline immunogenicity [2–5]. Systematic reviews of immune checkpoint inhibitors in MSS CRC continue to show limited efficacy for checkpoint blockade alone, while emphasizing that combination strategies will likely be required to improve outcomes in this population [5].

A key challenge is that MSS CRC is often described as “immune cold,” but this term can obscure substantial biological heterogeneity. Some MSS tumours may be true immune-desert tumours, characterized by low cytotoxic T-cell activity and weak IFNγ-associated chemokine recruitment. Others may show stromal-barrier or immune-excluded features, including cancer-associated fibroblast activity, extracellular-matrix remodelling, angiogenesis, hypoxia and suppressive myeloid programmes. A further subset may display partial immune activation together with checkpoint or suppressive counter-regulation. These distinct tumour states may require different therapeutic strategies. For example, immune-desert tumours may require immune priming through innate immune activation, radiotherapy, chemotherapy, vaccines or oncolytic viruses, whereas barrier-high tumours may require stromal, angiogenic, myeloid or TGFβ-associated remodelling before checkpoint blockade can be effective [5–7].

Several tumour-intrinsic and tumour-microenvironmental pathways have been implicated in MSS CRC immune resistance. WNT/β-catenin and MAPK signalling can contribute to immune exclusion or impaired antigen presentation. VEGF and hypoxia can suppress dendritic-cell and T-cell function while promoting abnormal vasculature and myeloid suppression. Cancer-associated fibroblasts and extracellular-matrix programmes can restrict immune infiltration and create an immunosuppressive stromal niche. Epigenetic regulators, DNA-damage response pathways and apoptosis-regulatory programmes may also modulate tumour-cell sensitivity to immune or cytotoxic stress [6–8]. These observations suggest that converting MSS CRC from an immune-cold to an immune-inflamed state will require rational combinations that address both tumour-cell-intrinsic vulnerabilities and microenvironmental barriers.

Large-scale functional genomics resources now provide an opportunity to complement patient transcriptomic analyses with tumour-cell dependency data. The Cancer Dependency Map (DepMap) project provides open-access genome-scale CRISPR and other functional-screening data across hundreds of cancer cell-line models, with the aim of identifying cancer vulnerabilities and supporting therapeutic discovery [9]. DepMap CRISPR gene-effect scores estimate the effect of gene knockout on cell growth or viability, with more negative scores indicating stronger dependency. Such data can help distinguish genes that are merely differentially expressed in patient tumours from genes that may represent tumour-cell-intrinsic vulnerabilities. At the same time, cell-line dependency data cannot fully capture stromal, immune or vascular dependencies, making integration with patient tumour transcriptomics essential for interpretation.

Integrating patient-derived immune-state signatures with cancer dependency data is therefore a promising strategy for target prioritization in MSS CRC. Patient transcriptomics can identify immune-cold, inflamed and barrier-high tumour states, while DepMap can highlight which candidate genes are functionally required in CRC tumour-cell models. This combined approach is particularly useful because MSS CRC resistance is unlikely to be explained by a single mechanism. Instead, actionable candidates may fall into at least two broad categories: tumour-cell-intrinsic vulnerabilities, such as receptor tyrosine kinase, PI3K, DNA-damage response, epigenetic or apoptosis-regulatory pathways; and patient-derived microenvironmental programmes, such as CAF/ECM, VEGF/hypoxia or myeloid-suppressive signatures. Separating these categories is necessary to avoid overinterpreting stromal markers as cancer-cell dependencies or overlooking druggable tumour-intrinsic pathways that may support combination therapy.

Several clinically relevant examples support this integrative logic. ERBB2/HER2 represents an established actionable target in a molecularly selected subset of metastatic CRC, and HER2-directed therapies have become increasingly relevant for patients with HER2-amplified or HER2-overexpressing disease [10,11]. VEGF-directed therapy is already part of CRC treatment and may also influence the immune microenvironment through effects on vascular normalization, hypoxia and immune suppression [7,8]. DNA-damage checkpoint targets such as ATR, WEE1 and CHEK1, epigenetic regulators such as HDACs and BET proteins, and apoptosis-regulatory proteins such as BCL2L1 and MCL1 are also of translational interest because they may increase tumour-cell stress, enhance antigenicity or sensitize tumours to cytotoxic and immune-mediated killing. However, these candidates need to be prioritized within the biological context of MSS immune-cold or barrier-high disease rather than considered as generic anticancer targets.

In this study, we integrated TCGA COAD/READ patient transcriptomics, MSIsensor-defined MSS/MSI-L classification, curated immune and stromal module scoring, focused differential expression, and DepMap CRISPR dependency data to prioritize candidate vulnerabilities in immune-cold MSS CRC. We first restricted the analysis to MSS/MSI-L tumours and classified them into immune-cold, hot/inflamed, barrier-high and intermediate transcriptomic states. We then compared MSS immune-cold tumours with MSS hot/inflamed tumours and MSS barrier-high tumours with MSS immune-cold tumours to identify cold- and barrier-associated programmes. Finally, we integrated patient-expression evidence with colorectal cancer DepMap dependency scores, immune relevance and druggability annotation to generate a ranked list of candidate targets and pathways. This approach is intended as a reproducible, hypothesis-generating framework that nominates tumour-cell-intrinsic vulnerabilities and microenvironmental barrier programmes for future validation, rather than as a clinically validated classifier or evidence of therapeutic efficacy.

## Results

### MSS/MSI-L colorectal cancers retain substantial immune-state heterogeneity

To focus specifically on the immunotherapy-resistant colorectal cancer population, we restricted the analysis to TCGA COAD/READ tumours classified as MSS/MSI-L using MSIsensor score <3.5. Among tumours with available RNA-seq expression data and usable MSIsensor scores, 494 MSS/MSI-L colorectal cancers were included in the analysis.

Using curated immune, stromal, tumour-intrinsic and suppressive gene-expression modules, MSS/MSI-L tumours were classified into four transcriptomic states: MSS immune-cold, MSS hot/inflamed, MSS barrier-high and MSS intermediate. The largest group was the MSS intermediate state, comprising 218 of 494 tumours, corresponding to 44.1% of the MSS/MSI-L cohort. MSS immune-cold tumours accounted for 102 cases, or 20.6%. MSS hot/inflamed tumours accounted for 91 cases, or 18.4%, while MSS barrier-high tumours accounted for 83 cases, or 16.8% (Fig 1a) (Table 1).

**Figure 1.**
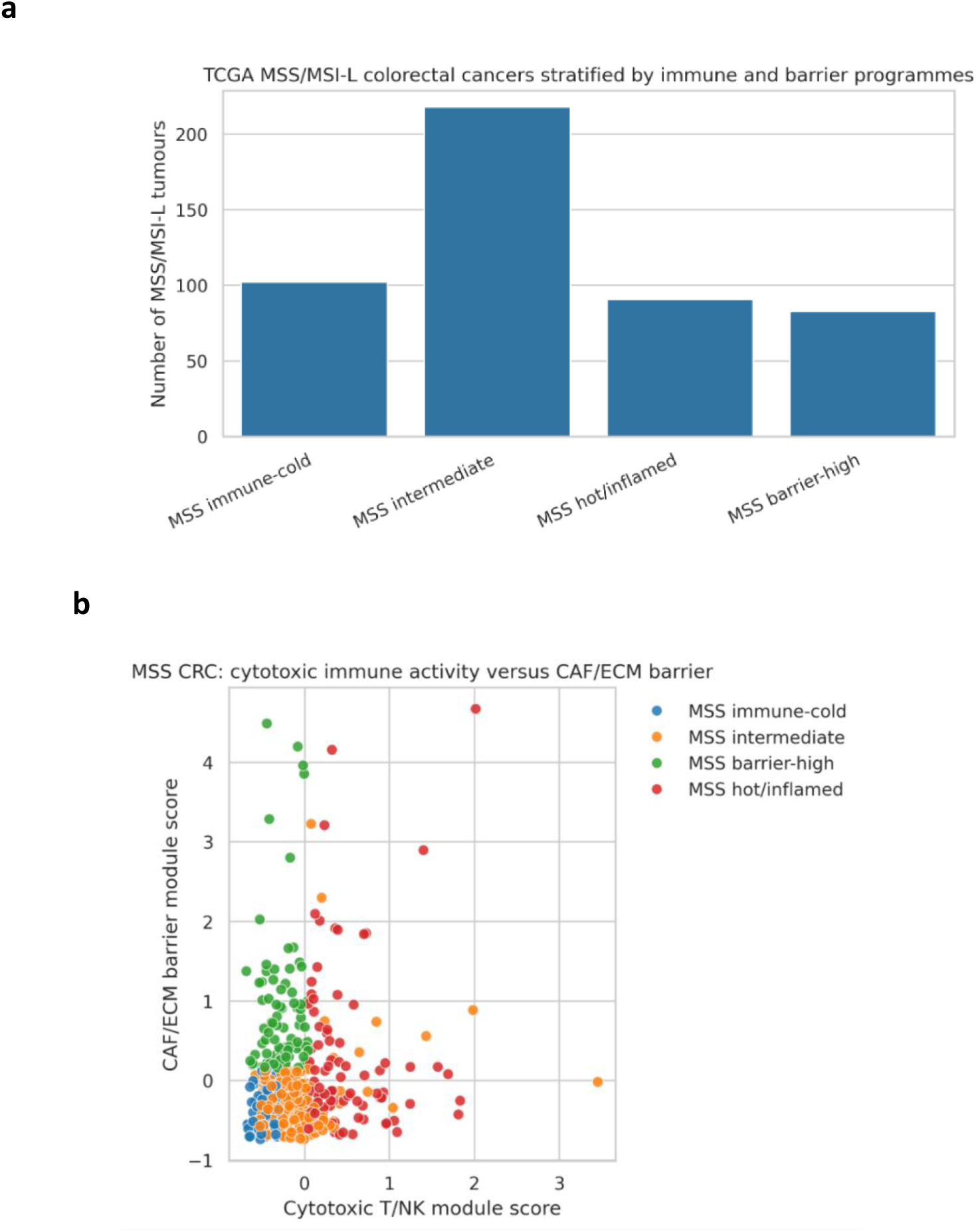
MSS/MSI-L colorectal cancers display heterogeneous transcriptomic immune and stromal states: TCGA COAD/READ tumours with available RNA-seq data and numeric MSIsensor scores were restricted to the MSS/MSI-L subgroup using MSIsensor score <3.5. (A) Distribution of MSS/MSI-L transcriptomic states among 494 tumours. The largest subgroup was MSS intermediate (218/494; 44.1%), followed by MSS immune-cold (102/494; 20.6%), MSS hot/inflamed (91/494; 18.4%) and MSS barrier-high (83/494; 16.8%). (B) Scatter plot of cytotoxic T/NK module score versus CAF/ECM barrier module score. Each point represents one MSS/MSI-L tumour and is coloured by transcriptomic state. The plot illustrates separation of tumours along immune-activation and stromal-barrier axes.

**Table 1.** Distribution of transcriptomic states among MSIsensor-defined MSS/MSI-L TCGA colorectal tumours: Tumours were restricted to the MSS/MSI-L subgroup using MSIsensor score <3.5 and classified into MSS intermediate, MSS immune-cold, MSS hot/inflamed and MSS barrier-high states using curated immune and stromal module scores.

| *MSS/MSI-L transcriptomic state* | *Number of tumours* | *Percentage* |
| --- | --- | --- |
| *MSS intermediate* | 218 | 44.1% |
| *MSS immune-cold* | 102 | 20.6% |
| *MSS hot/inflamed* | 91 | 18.4% |
| *MSS barrier-high* | 83 | 16.8% |
| *Total* | 494 | 100% |

These findings indicate that MSS/MSI-L colorectal cancer is not uniformly immune-cold. Instead, even within the MSS/MSI-L subgroup, tumours display heterogeneous immune and stromal states, including a subset with inflamed immune features and a subset with stromal-barrier-high biology.

### MSS immune-cold tumours are depleted for cytotoxic, IFNγ-chemokine and adaptive immune engagement programmes

We next compared MSS immune-cold tumours with MSS hot/inflamed tumours to define immune programmes associated with the cold MSS phenotype. Focused gene-level comparison showed that MSS immune-cold tumours were strongly depleted for canonical cytotoxic and inflammatory immune genes relative to MSS hot/inflamed tumours (Fig 1b).

Among the most significantly reduced genes in MSS immune-cold tumours were NKG7, CXCL10, CD8A, CXCL9 and LAG3 (Table 2). NKG7 expression was markedly lower in MSS immune-cold tumours than in MSS hot/inflamed tumours, consistent with reduced cytotoxic lymphocyte activity. Similarly, CD8A was reduced in MSS immune-cold tumours, indicating lower CD8 T-cell infiltration or activity. IFNγ-associated chemokines, including CXCL9 and CXCL10, were also substantially reduced, suggesting impaired effector T-cell recruitment. LAG3 was also lower in MSS immune-cold tumours, consistent with reduced adaptive immune engagement rather than active checkpoint-mediated restraint.

**Table 2.** Representative genes depleted in MSS immune-cold tumours relative to MSS hot/inflamed tumours: Focused gene-level comparison was performed using Mann–Whitney U tests with Benjamini–Hochberg correction. The table lists representative cytotoxic, chemokine and adaptive immune engagement genes with lower expression in MSS immune-cold tumours.

| *Gene* | *Mean expression in MSS immune-cold* | *Mean expression in MSS hot/inflamed* | *Direction* |
| --- | --- | --- | --- |
| *NKG7* | 22.93 | 175.92 | Lower in cold |
| *CXCL10* | 71.06 | 1120.07 | Lower in cold |
| *CD8A* | 34.01 | 260.88 | Lower in cold |
| *CXCL9* | 108.71 | 1619.74 | Lower in cold |
| *LAG3* | 23.20 | 134.50 | Lower in cold |

Together, these findings support the classification of the MSS immune-cold state as a tumour phenotype with weak cytotoxic infiltration, reduced inflammatory chemokine signalling and limited adaptive immune activation (Fig 2 a-d).

**Figure 2.**
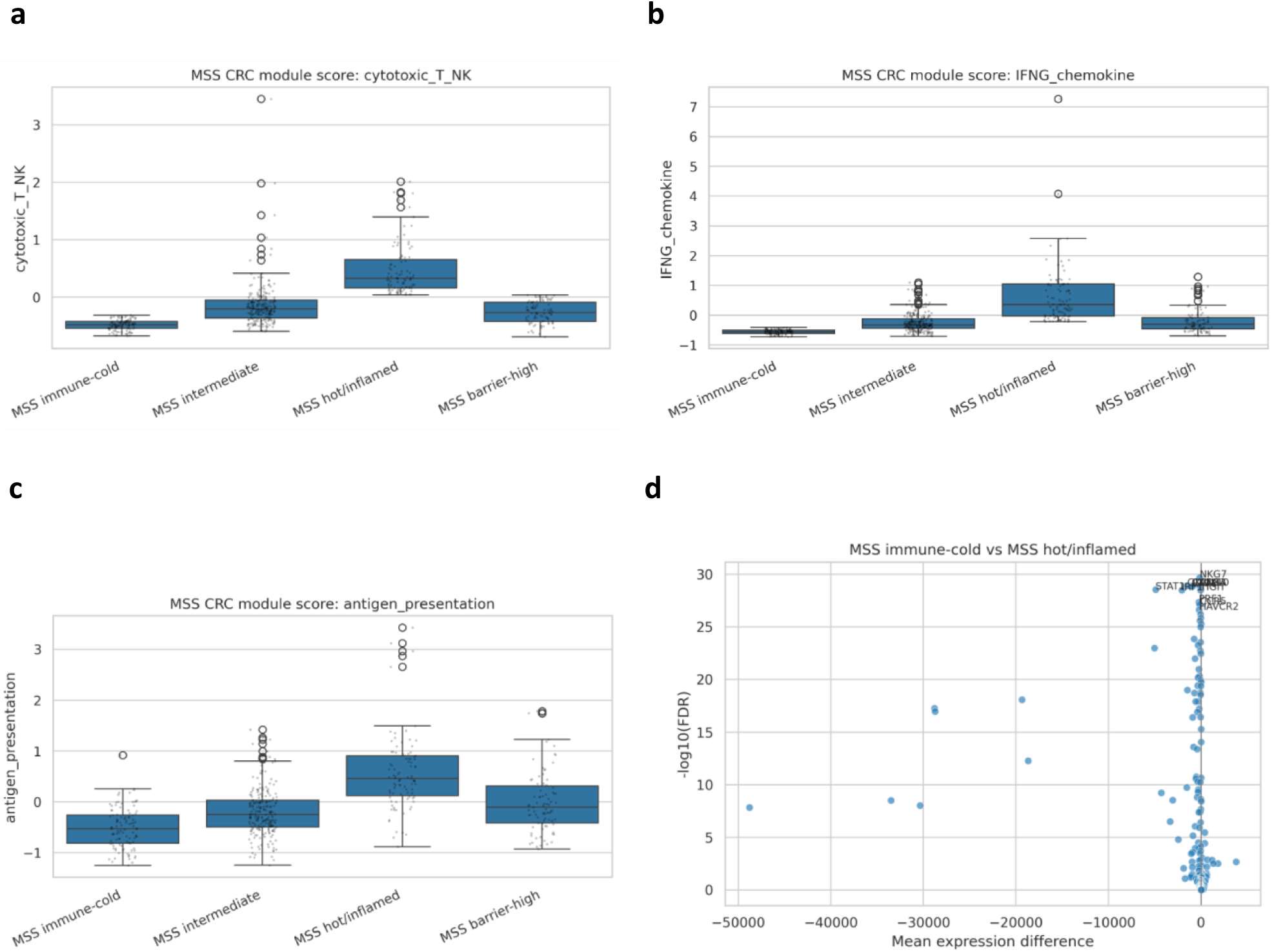
MSS immune-cold tumours are depleted for cytotoxic and inflammatory immune programmes: Curated module scores were calculated as the mean z-scored expression of available genes within each module. (A) Cytotoxic T/NK module scores across MSS/MSI-L transcriptomic states. (B) IFNγ/chemokine module scores across MSS/MSI-L transcriptomic states. (C) Antigen-presentation module scores across MSS/MSI-L transcriptomic states. (D) Focused gene-level comparison of MSS immune-cold versus MSS hot/inflamed tumours. MSS immune-cold tumours were depleted for canonical immune-effector and inflammatory genes, including NKG7, CXCL10, CD8A, CXCL9 and LAG3. Gene-level statistical testing was performed using Mann–Whitney U tests with Benjamini–Hochberg false-discovery-rate correction.

### MSS barrier-high tumours show stromal and extracellular-matrix enrichment

We next compared MSS barrier-high tumours with MSS immune-cold tumours to identify features associated with stromal-barrier biology. MSS barrier-high tumours showed increased CAF/ECM-associated expression, consistent with a stromal-rich and putatively immune-restrictive phenotype (Fig 3a).

**Figure 3.**
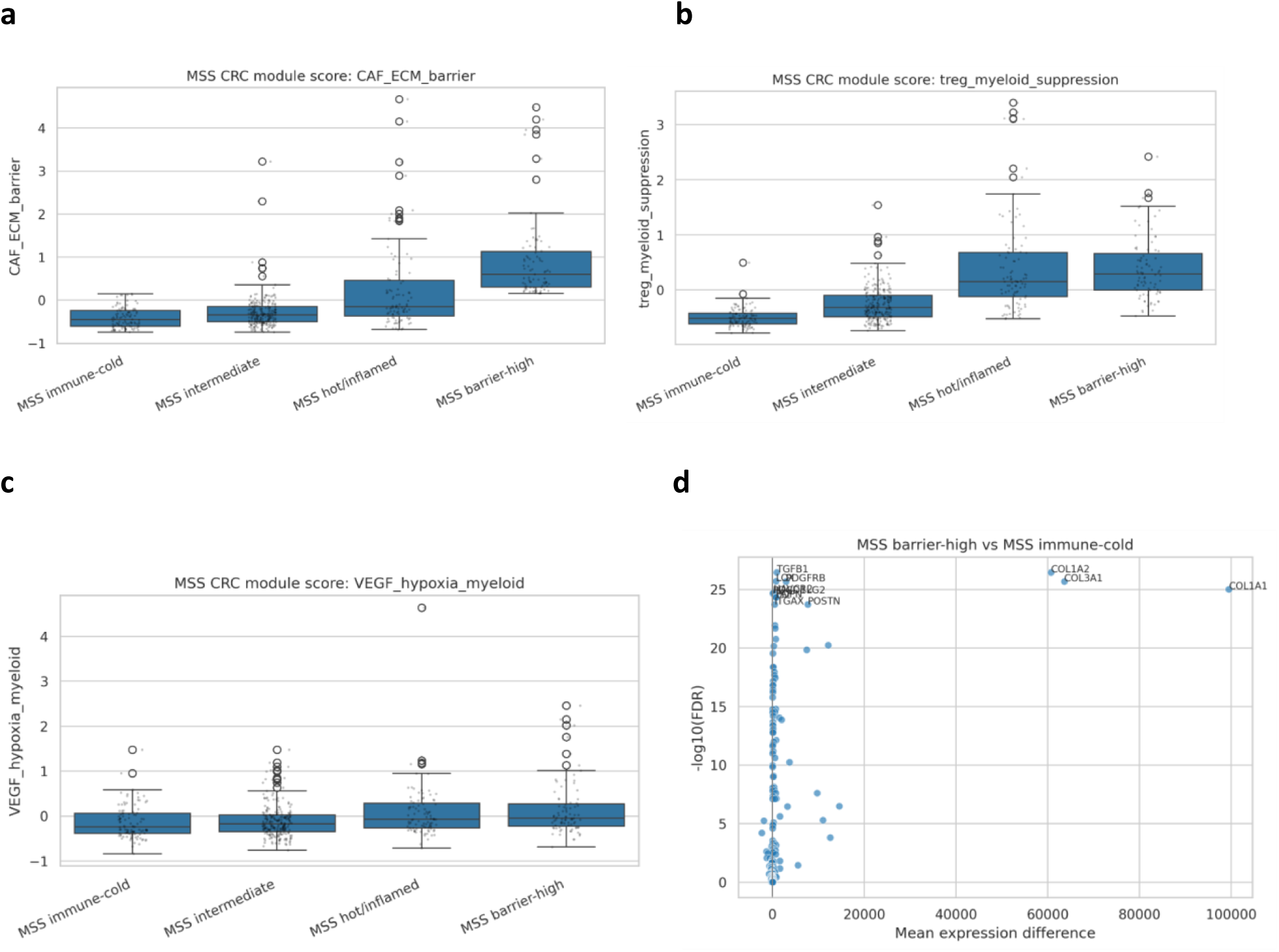
MSS barrier-high tumours are enriched for stromal and suppressive microenvironmental programmes: Module-score distributions were compared across MSS/MSI-L transcriptomic states. (A) CAF/ECM barrier module scores were increased in MSS barrier-high tumours. (B) Treg/myeloid suppression module scores across MSS states. (C) VEGF/hypoxia/myeloid module scores across MSS states. (D) Focused gene-level comparison of MSS barrier-high versus MSS immune-cold tumours. Barrier-high tumours showed enrichment of extracellular-matrix and stromal-associated genes, including COL1A1, COL1A2 and COL3A1. These genes were interpreted as patient-derived stromal-barrier markers rather than tumour-cell-intrinsic dependencies. Statistical comparisons were performed using Mann–Whitney U tests with Benjamini–Hochberg correction.

The integrated ranking and gene-level analysis identified strong barrier-associated expression of extracellular-matrix genes, including COL1A1, COL1A2 and COL3A1 (Fig 3d). These genes showed prominent elevation in the barrier-high state and represented the dominant patient-derived stromal signature. Because collagen and ECM genes are primarily microenvironmental markers rather than tumour-cell-intrinsic dependencies, these findings were interpreted as evidence of a barrier-associated patient phenotype rather than as direct DepMap-supported tumour dependencies.

The barrier-high phenotype therefore represents a biologically distinct MSS/MSI-L state characterized by extracellular-matrix remodelling and stromal enrichment. This state may require therapeutic strategies directed toward stromal remodelling, angiogenic suppression, CAF/TGFβ-associated pathways or myeloid reprogramming rather than immune priming alone (Fig 3b-c).

### Module-level differential analysis distinguishes cold, hot and barrier-high MSS colorectal cancer states

To summarize pathway-level differences between MSS/MSI-L states, we performed module-level differential testing. The analysis compared MSS immune-cold versus MSS hot/inflamed and MSS barrier-high versus MSS immune-cold tumours (Fig 4).

**Figure 4.**
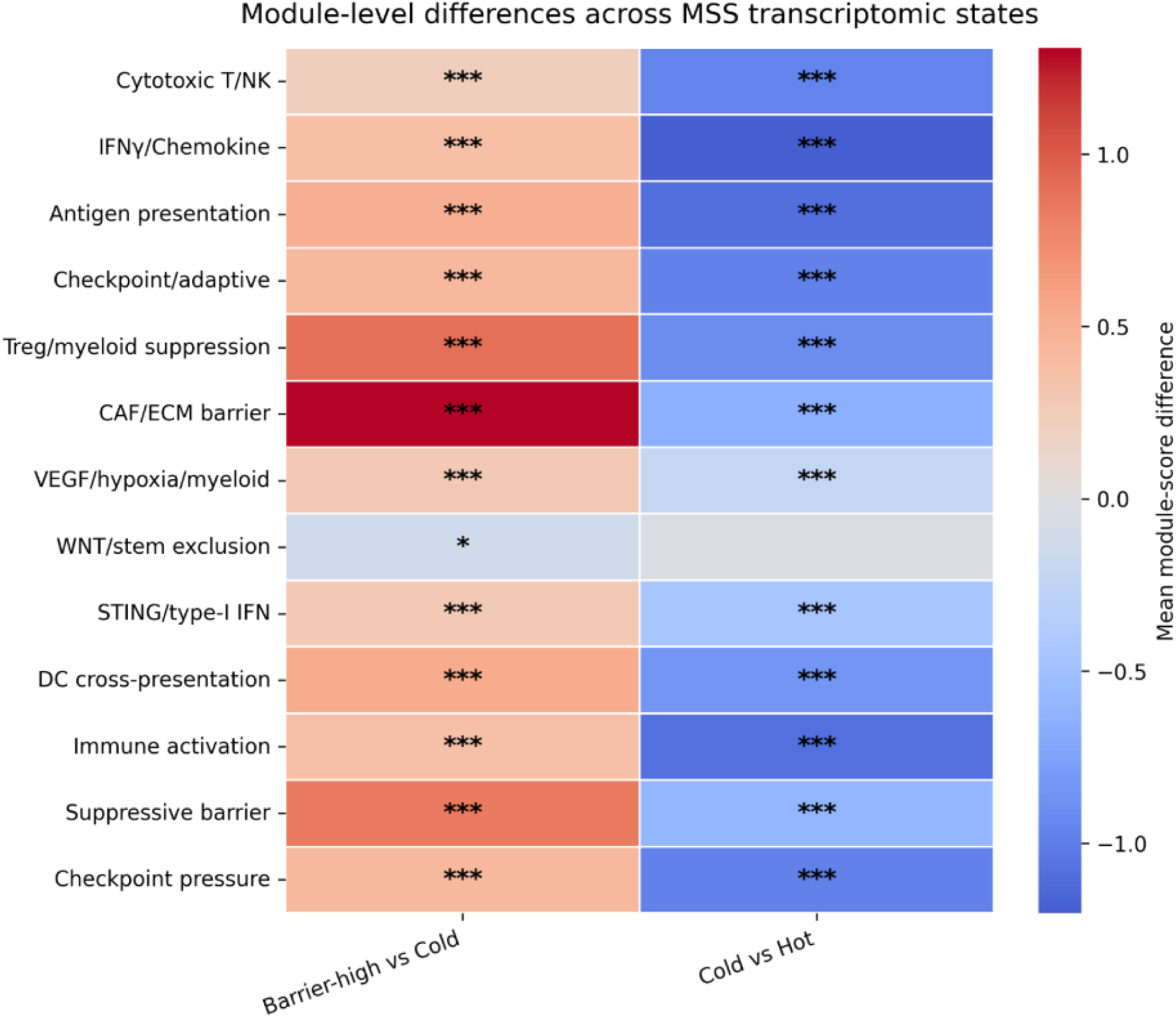
Module-level differential analysis distinguishes MSS immune-cold, hot/inflamed and barrier-high states: Heatmap showing mean module-score differences for the main comparisons: MSS immune-cold versus MSS hot/inflamed and MSS barrier-high versus MSS immune-cold. MSS immune-cold tumours showed reduced cytotoxic T/NK, IFNγ/chemokine and antigen-presentation activity relative to MSS hot/inflamed tumours. MSS barrier-high tumours showed increased CAF/ECM barrier and associated suppressive microenvironmental features relative to MSS immune-cold tumours. Asterisks indicate FDR-adjusted significance from module-level comparisons.

MSS immune-cold tumours showed reduced cytotoxic T/NK, IFNγ-chemokine and antigen-presentation activity relative to MSS hot/inflamed tumours (Fig 4). This confirmed that the cold phenotype was not limited to one marker gene but reflected coordinated depletion of immune-effector and inflammatory recruitment programmes.

By contrast, MSS barrier-high tumours were characterized by increased CAF/ECM barrier activity and associated suppressive microenvironmental features relative to MSS immune-cold tumours (Fig 4). These results support a model in which MSS/MSI-L tumours can fail immune control through at least two distinct mechanisms: poor immune activation in the immune-cold state and stromal-barrier/suppressive microenvironment enrichment in the barrier-high state.

### DepMap CRISPR screening identifies tumour-cell-intrinsic dependencies among candidate targets

To distinguish patient-derived immune or stromal signatures from tumour-cell-intrinsic vulnerabilities, we integrated the MSS patient transcriptomic analysis with DepMap CRISPR gene-effect data. DepMap dependency data were available for 1208 total cancer models, including 63 colorectal cancer models.

Several strong colorectal cancer dependencies were identified among the candidate gene set. As expected, core proliferative and mitotic regulators such as AURKB, AURKA, CDK1 and TOP2A showed strong dependency evidence in colorectal cancer cell lines (Fig 5a). For example, AURKB showed a mean gene effect of −2.12 in colorectal cancer models, indicating a strong growth defect following gene knockout. AURKA and ATR also showed substantial dependency evidence, with mean colorectal cancer gene-effect values of −1.18 and −0.96, respectively (Table 3).

**Figure 5.**
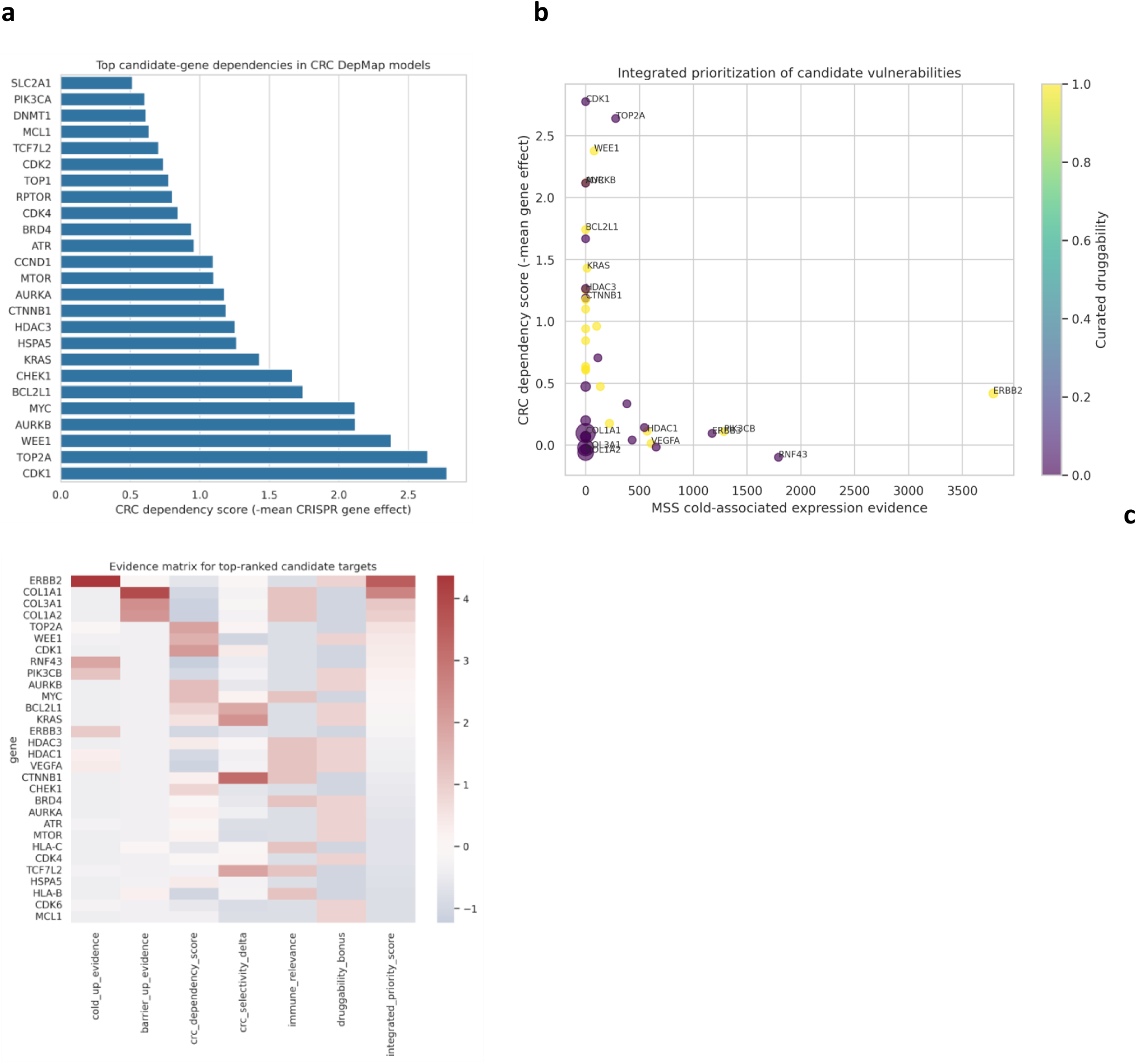
DepMap and integrated prioritization nominate tumour-cell-intrinsic and clinically actionable candidate vulnerabilities: DepMap CRISPR gene-effect data were summarized across 1208 total cancer models, including 63 colorectal cancer models. More negative gene-effect values indicate stronger growth impairment after gene knockout. (A) Representative top colorectal cancer dependencies among candidate genes. Strong dependencies included broad mitotic or DNA-damage response genes such as AURKB, AURKA and ATR, which were interpreted cautiously because of potential common essentiality. (B) Integrated candidate-prioritization bubble plot combining patient-expression evidence, CRC dependency score and curated druggability. (C) Evidence matrix for top-ranked candidate targets.

**Figure 6.**
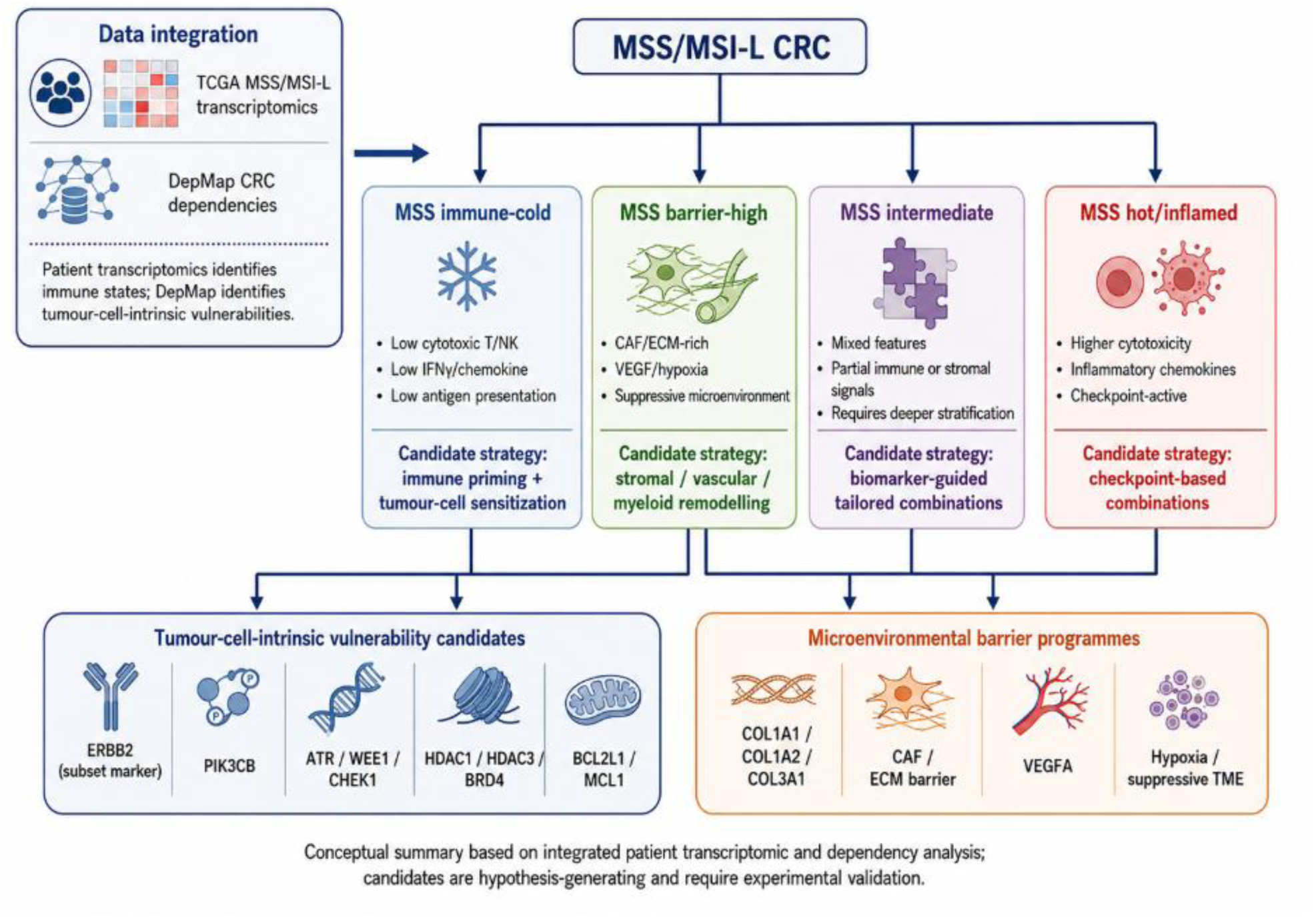
Integrated framework for state-matched candidate therapeutic strategies in MSS colorectal cancer: Schematic model summarizing the proposed state-matched framework. MSS immune-cold tumours show low cytotoxic T/NK, IFNγ-chemokine and antigen-presentation activity and may require immune priming together with tumour-cell sensitization. MSS barrier-high tumours show CAF/ECM, angiogenic/hypoxic and suppressive microenvironmental programmes and may require stromal, vascular or myeloid remodelling. MSS hot/inflamed tumours retain cytotoxic and inflammatory immune features and may be more aligned with checkpoint-based combinations. MSS intermediate tumours may require further molecular stratification. Candidate tumour-cell-intrinsic vulnerabilities include ERBB2, PIK3CB, ATR/WEE1/CHEK1, HDAC/BRD4 and apoptosis-regulatory genes, whereas COL1A1, COL1A2 and COL3A1 represent stromal-barrier signatures rather than tumour-cell dependencies.

**Table 3.** Representative candidate-gene dependencies in DepMap colorectal cancer models: DepMap CRISPR gene-effect scores were summarized across colorectal cancer cell-line models. More negative gene-effect values indicate stronger growth impairment after gene knockout. Broadly essential genes were interpreted cautiously and distinguished from immune-state-specific candidate vulnerabilities.

| *Gene* | *Mean gene effect in CRC models* | *Interpretation* |
| --- | --- | --- |
| AURKB | -2.12 | Strong dependency, broad mitotic essentiality |
| *AURKA* | -1.18 | Strong dependency, mitotic regulator |
| *ATR* | -0.96 | DNA-damage response dependency |
| *APC* | -0.40 | Moderate CRC-relevant dependency |
| *B2M* | +0.03 | Not a tumour-cell dependency |
| *ACTA2* | +0.04 | Stromal marker, not a tumour-cell dependency |

However, many of these genes also represent broad essential genes across cancer types and were therefore interpreted cautiously. Such genes may represent tumour-cell-intrinsic sensitization candidates but are less likely to be immune-state-specific targets. The analysis therefore incorporated a common-essentiality penalty to reduce over-prioritization of broadly essential cell-cycle genes.

As expected, stromal and immune microenvironment markers such as ACTA2, ARG1 and B2M did not behave as strong tumour-cell-line dependencies in DepMap. This distinction reinforces the complementary nature of the analysis: patient transcriptomics identifies tumour microenvironment-associated states, whereas DepMap primarily identifies tumour-cell-intrinsic vulnerabilities.

### Integrated prioritization highlights clinically actionable, druggable and immune-relevant candidates for MSS immune-cold colorectal cancer

We next integrated patient transcriptomic evidence, DepMap dependency evidence, immune relevance and druggability annotation to prioritize candidate targets for MSS immune-cold and barrier-associated colorectal cancer (Fig 5b, c). The integrated ranking highlighted several candidate classes, including clinically actionable receptor tyrosine kinases, angiogenic and stromal-associated pathways, PI3K signalling, DNA-damage checkpoint dependencies, epigenetic regulators and apoptosis-sensitization targets.

ERBB2 emerged as the highest-ranked clinically actionable candidate signal in the integrated analysis (Fig 5c). In focused gene-level testing, ERBB2 expression was significantly higher in MSS immune-cold tumours than in MSS hot/inflamed tumours, with mean expression values of 8457.04 and 4668.65, respectively (Supplementary fig 2). The mean difference was 3788.39, with an FDR-adjusted P value of 0.00205. ERBB2 expression was further increased in MSS barrier-high tumours compared with MSS immune-cold tumours, with mean expression values of 13986.29 and 8457.04, respectively. The mean difference was 5529.25, with an FDR-adjusted P value of 0.03547 (Supplementary fig 2). In DepMap colorectal cancer models, ERBB2 showed modest dependency evidence, supporting its nomination as a candidate actionable subset-associated signal rather than as a universal MSS immune-cold dependency. Because HER2-directed treatment selection in CRC is based on amplification, immunohistochemistry, *in situ* hybridization or sequencing-based criteria, ERBB2 mRNA elevation should be considered a prioritization signal requiring orthogonal molecular validation.

Other clinically or therapeutically relevant candidates included VEGFA, PIK3CB, ATR, WEE1, HDAC1, HDAC3, BRD4, BCL2L1 and MCL1. VEGFA was prioritized as an immune-relevant and clinically actionable angiogenic axis, consistent with the role of vascular and hypoxia-associated programmes in immune suppression (Table 4). PIK3CB represented a PI3K-pathway candidate associated with immune-cold features. ATR, WEE1 and CHEK1 represented DNA-damage checkpoint vulnerabilities that may be relevant for immunogenic stress or combination sensitization strategies. HDAC1, HDAC3 and BRD4 represented epigenetic candidates with potential immune-modulatory relevance. BCL2L1 and MCL1 represented apoptosis-priming candidates that may influence tumour-cell sensitivity to cytotoxic stress.

**Table 4.** Representative candidate targets nominated by integrated patient transcriptomic and DepMap dependency analysis: Candidate genes were prioritized using patient expression evidence, CRC dependency evidence, immune relevance, druggability annotation and common-essentiality penalty. The table summarizes candidate target classes including ERBB2, VEGFA, PI3K, DNA-damage response, epigenetic and apoptosis-regulatory pathways, as well as stromal-barrier marker genes.

| *Candidate* | *Main evidence* | *Interpretation* |
| --- | --- | --- |
| *ERBB2* | Cold-up, barrier-up, clinically actionable, modest CRC dependency | Candidate actionable subset marker |
| *VEGFA* | Cold-up, immune-relevant, clinically actionable | Angiogenic/immune-suppressive axis |
| *PIK3CB* | Cold-up, druggable pathway | PI3K-pathway candidate |
| *ATR / WEE1 / CHEK1* | CRC dependency, druggable pathway | DNA-damage checkpoint vulnerabilities |
| *HDAC1 / HDAC3 / BRD4* | Druggable, immune-relevant epigenetic regulators | Candidate immune-modulatory targets |
| *BCL2L1 / MCL1* | CRC dependency, druggable apoptosis pathway | Apoptosis-sensitization candidates |
| *COL1A1 / COL1A2 / COL3A1* | Strong barrier-high patient signature | Stromal-barrier markers, not tumour-cell dependencies |

In parallel, collagen genes including COL1A1, COL1A2 and COL3A1 ranked highly because of strong barrier-associated patient expression. These genes were interpreted as markers of the stromal-barrier phenotype rather than direct tumour-cell-intrinsic vulnerabilities. Their presence in the integrated ranking supports the importance of distinguishing druggable tumour-cell dependencies from microenvironmental barrier programmes.

### Patient–dependency integration separates tumour-cell-intrinsic vulnerabilities from microenvironmental barrier programmes

The integrated analysis revealed two complementary classes of candidate vulnerabilities. The first class consisted of tumour-cell-intrinsic dependencies captured by colorectal cancer DepMap models. These included DNA-damage response genes, mitotic regulators, apoptosis regulators, epigenetic regulators and selected clinically actionable oncogenic pathways. These candidates may be most relevant for tumour-cell sensitization strategies, particularly when combined with immune priming or checkpoint-based therapy.

The second class consisted of patient-derived microenvironmental programmes, particularly CAF/ECM and collagen-rich barrier signatures. These were strongly represented in MSS barrier-high tumours but were not captured as tumour-cell dependencies in DepMap. This distinction is biologically important because stromal-barrier biology is unlikely to be fully modelled by cancer cell-line screens. Instead, such features may require therapeutic strategies targeting the tumour microenvironment, including stromal remodelling, anti-angiogenic therapy, CAF/TGFβ-associated approaches or myeloid reprogramming.

Together, these findings support a hypothesis-generating, state-matched therapeutic framework for MSS/MSI-L colorectal cancer. MSS immune-cold tumours may require immune priming and tumour-cell sensitization. MSS barrier-high tumours may require stromal or suppressive microenvironment remodelling. MSS hot/inflamed tumours may be more biologically aligned with checkpoint-based therapeutic strategies. MSS intermediate tumours may require additional stratification based on dominant tumour-intrinsic or microenvironmental vulnerabilities.

### Summary of main finding

MSS/MSI-L CRC contains distinct immune-cold, hot, intermediate and barrier-high states. Patient transcriptomics and DepMap integration separate stromal-barrier programmes from tumour-cell-intrinsic dependencies and nominate actionable candidate pathways for future immune-conversion strategies.

## Discussion

In this study, TCGA patient transcriptomics, MSIsensor-defined MSS/MSI-L classification, curated immune and stromal module scores, focused gene-level comparisons and DepMap CRISPR dependency data were combined to prioritize candidate vulnerabilities in immune-cold MSS colorectal cancer. The central observation was that MSS/MSI-L CRC is not a single immune-cold entity. Among 494 MSS/MSI-L tumours, 218 were classified as MSS intermediate, 102 as MSS immune-cold, 91 as MSS hot/inflamed and 83 as MSS barrier-high. This finding is important because MSS CRC is often considered as one immunotherapy-resistant group, whereas the present analysis indicates that distinct immune and stromal states remain detectable within MSS/MSI-L disease.

The MSS immune-cold state was characterized by depletion of cytotoxic, IFNγ-chemokine and adaptive immune engagement programmes relative to MSS hot/inflamed tumours. At the gene level, MSS immune-cold tumours showed markedly lower expression of NKG7, CD8A, CXCL9, CXCL10 and LAG3. These genes represent cytotoxic lymphocyte activity, CD8 T-cell infiltration, IFNγ-associated effector-cell recruitment and adaptive immune activation. The biological implication is that at least a subset of MSS CRC lacks the baseline immune contexture required for checkpoint blockade to be effective. This agrees with current clinical and translational understanding: checkpoint inhibitors have limited activity in most MSS/pMMR CRCs, and resistance is increasingly viewed as a failure of immune priming, trafficking, infiltration, antigen presentation and local immune activation rather than only a failure of checkpoint release [4,5,14].

A useful aspect of the analysis is the separation of immune-cold MSS tumours from barrier-high MSS tumours. MSS barrier-high tumours were enriched for extracellular-matrix and stromal features, including strong patient-derived signals from COL1A1, COL1A2 and COL3A1. These findings suggest that some MSS tumours may not be simple immune-desert tumours but may instead display a stromal-barrier or suppressive microenvironmental phenotype. This distinction has therapeutic relevance. Immune-cold tumours with low cytotoxic and chemokine activity may require immune-priming strategies such as radiotherapy, immunogenic chemotherapy, STING/TLR activation, vaccines or oncolytic viruses. By contrast, barrier-high tumours may require microenvironmental remodelling, including anti-angiogenic therapy, CAF/ECM targeting, TGFβ-associated approaches or myeloid reprogramming. Recent reviews of MSS CRC immunotherapy resistance similarly emphasize that resistance is conditional and multi-layered, involving both tumour-cell-intrinsic and microenvironmental mechanisms [5–8,14,15].

The integration with DepMap CRISPR dependency data adds a second layer of biological interpretation. Patient transcriptomics captures both tumour-cell and microenvironmental signals, whereas DepMap primarily captures tumour-cell-intrinsic dependencies in cancer cell-line models. This distinction is critical. In our analysis, stromal genes such as ACTA2 and immune or antigen-presentation genes such as B2M were not strong tumour-cell dependencies, as expected. Conversely, mitotic and DNA-damage response genes such as AURKB, AURKA and ATR showed strong colorectal cancer dependency evidence. DepMap guidance and analyses emphasize that more negative gene-effect scores indicate stronger dependency, but also that common essential genes should not be interpreted as tumour-specific vulnerabilities without caution [9,12,13,16,17,20]. We therefore incorporated a common-essentiality penalty and interpreted broad cell-cycle dependencies as potential sensitization candidates rather than immune-state-specific targets.

The integrated prioritization highlighted several translationally relevant candidate classes. ERBB2 was the highest-ranked clinically actionable candidate signal. ERBB2 expression was higher in MSS immune-cold tumours than in MSS hot/inflamed tumours and was further increased in MSS barrier-high tumours. Importantly, ERBB2 showed only modest DepMap dependency evidence; therefore, this finding should not be interpreted as evidence that all MSS immune-cold tumours are HER2-dependent. Rather, ERBB2 may mark a molecularly selected actionable subset within immune-cold or barrier-high MSS CRC. This is clinically plausible because HER2 amplification or overexpression is an established therapeutic target in a subset of metastatic CRC, and recent reviews continue to highlight HER2-directed therapy as an expanding precision-oncology strategy in colorectal cancer [10,11]. ERBB2 is therefore best framed as a candidate subset marker and pathway for further validation, not as a universal MSS immune-cold dependency.

VEGFA also emerged as an immune-relevant and clinically actionable candidate. This is biologically consistent with the role of VEGF in abnormal tumour vasculature, hypoxia, myeloid recruitment, dendritic-cell dysfunction and impaired T-cell infiltration. Anti-VEGF therapy is already used in CRC, and its potential immunomodulatory effects provide a rationale for combination approaches in selected MSS tumours. In the context of our analysis, VEGFA is especially relevant because it bridges patient-derived immune suppression and clinically actionable therapy. It may be particularly important in tumours with barrier-high, hypoxic or myeloid-suppressive features rather than in all MSS CRCs uniformly.

The DNA-damage response candidates ATR, WEE1 and CHEK1 are another important class emerging from the integrated framework. ATR showed strong CRC dependency evidence in DepMap, while WEE1 and CHEK1 represent druggable checkpoint kinases involved in replication stress and cell-cycle control. DNA-damage response inhibition is of interest in CRC because it may increase tumour-cell stress, enhance genomic instability, interact with chemotherapy or radiation, and potentially increase immunogenicity under appropriate conditions [18,19]. However, these agents are not intrinsically immune-specific, and their value in immune-cold MSS CRC will likely depend on rational combinations, biomarker selection and tolerability. Therefore, DDR candidates should be presented as tumour-cell sensitization strategies that may complement immune priming rather than as stand-alone immune-conversion therapies.

Epigenetic candidates, including HDAC1, HDAC3 and BRD4, also warrant discussion. Epigenetic regulation can influence antigen presentation, interferon signalling, endogenous retroviral expression, inflammatory gene programmes and T-cell exclusion. HDAC and BET inhibition have been explored as immune-modulatory strategies in cancer, although effects can be context-dependent and toxicity remains a concern. In this analysis, these genes were prioritized because they combined druggability, immune relevance and dependency evidence. This supports their inclusion as candidate sensitization pathways for future experimental testing in MSS immune-cold CRC models.

The apoptosis-regulatory candidates BCL2L1 and MCL1 represent a further class of tumour-cell-intrinsic vulnerabilities. These genes may not directly convert a tumour from cold to hot, but they may influence susceptibility to cytotoxic stress, immune-mediated killing or combination therapy. Apoptosis-priming approaches could therefore be relevant in combination with chemotherapy, radiation, targeted therapy or immune-priming strategies. As with DDR and epigenetic candidates, the main value of these targets is not that they explain immune coldness alone, but that they may identify tumour-cell vulnerabilities that can be exploited alongside immune-conversion strategies.

The study also highlights a major conceptual point: patient-derived microenvironmental signals and cell-line-derived dependencies should not be combined. Collagen genes such as COL1A1, COL1A2 and COL3A1 ranked highly because of strong barrier-associated expression in patient tumours, not because they are tumour-cell-intrinsic dependencies in DepMap. This is not a weakness; but, it demonstrates the value of the integrative design. Cell-line screens are powerful for identifying tumour-cell vulnerabilities but cannot model the full tumour microenvironment, including CAFs, extracellular matrix, endothelial cells, macrophages or T cells. Therefore, stromal-barrier genes should be interpreted as markers of a patient phenotype that may require microenvironment-directed therapy, while DepMap-supported genes should be interpreted as tumour-cell-intrinsic vulnerability candidates.

The main translational implication is that MSS CRC may require state-matched combination strategies. MSS immune-cold tumours may need immune priming plus tumour-cell sensitization. MSS barrier-high tumours may require stromal, vascular or myeloid remodelling before immune activation can become effective. MSS hot/inflamed tumours, although still MSS/MSI-L, may be more biologically aligned with checkpoint-based combinations. MSS intermediate tumours may require deeper molecular stratification to determine whether tumour-intrinsic dependencies, stromal suppression, WNT/stemness, angiogenesis or myeloid programmes dominate. This framework may help avoid one of the common pitfalls in MSS CRC immunotherapy development: testing broad combinations in unselected MSS populations where multiple resistance mechanisms coexist.

Several strengths support the study. First, the analysis focuses specifically on MSS/MSI-L CRC, the subgroup with the greatest unmet need in immunotherapy. Second, the study integrates patient-level transcriptomic states with functional cancer dependency data, rather than relying only on differential expression. Third, the prioritization strategy explicitly separates tumour-cell-intrinsic dependency evidence from patient-derived stromal and immune signatures. Fourth, the final output is translationally interpretable, nominating candidate target classes including HER2/ERBB2, VEGF, PI3K, DNA-damage response, epigenetic regulation and apoptosis priming. This provides a practical hypothesis-generating resource for future experimental and clinical studies.

The study also has important limitations. First, the patient component was based on bulk RNA-seq, which cannot resolve cell-of-origin or spatial localization. Stromal-barrier features such as collagen expression may reflect CAF-rich or ECM-rich tumour microenvironments, but bulk transcriptomics cannot prove physical T-cell exclusion. Spatial transcriptomics, multiplex immunofluorescence or imaging mass cytometry would be required to validate whether barrier-high tumours truly restrict cytotoxic lymphocyte infiltration. Second, DepMap dependency data are derived from cancer cell lines and therefore do not contain stromal, immune, endothelial or extracellular-matrix dependencies. This limitation is particularly important for MSS barrier-high disease. Third, the integrated ranking includes curated druggability and immune-relevance annotations, which are useful for prioritization but inevitably involve some assumptions. Fourth, the state definitions are cohort-relative and partly based on the immune and stromal modules later compared between groups; therefore, the module-level results should be viewed as descriptive internal validation rather than as an independent discovery test. Fifth, the current analysis does not include independent cohort validation, survival or treatment-response modelling, consensus molecular subtype adjustment or formal tumour-purity correction. Finally, the analysis does not test whether candidate targets experimentally convert cold MSS CRC into inflamed tumours. Therefore, all candidates should be considered hypotheses for future validation, not confirmed therapeutic strategies. Future work should also integrate mutation status, copy-number alteration, consensus molecular subtype, tumour mutational burden, microbiome features, tumour purity, spatial immune architecture and treatment-response data.

Despite these limitations, the present analysis provides a useful translational framework. It suggests that immune-cold MSS CRC should not be approached as a single therapeutic category. Instead, therapeutic prioritization should distinguish immune-desert biology, stromal-barrier biology, tumour-cell-intrinsic dependency and clinically actionable molecular subsets. By integrating patient transcriptomics with DepMap dependency data, this study nominates candidate targets and pathways that can be tested in mechanistic models of MSS CRC immune resistance. The most immediate next step would be to validate top candidates such as ERBB2, VEGFA, ATR/WEE1/CHEK1, HDAC/BRD4 and BCL2L1/MCL1 in MSS CRC organoid–immune co-culture systems, with readouts including antigen presentation, IFNγ/CXCL9/CXCL10 induction, T-cell recruitment, tumour-cell killing and response to checkpoint blockade.

In conclusion, MSS/MSI-L colorectal cancer contains distinct immune-cold, hot/inflamed, barrier-high and intermediate transcriptomic states. Patient transcriptomics identified immune-depleted and stromal-barrier programmes, while DepMap integration prioritized tumour-cell-intrinsic dependencies and clinically actionable candidates. The resulting framework separates microenvironmental barrier signatures from cancer-cell vulnerabilities and supports state-matched strategies for future immune-conversion studies in MSS CRC.

## Materials and Methods

### Study design

This study was designed as a translational analysis of publicly available patient transcriptomic and cancer-dependency datasets to identify candidate vulnerabilities associated with immune-cold microsatellite-stable/microsatellite instability-low colorectal cancer. Patient tumour transcriptomic data from TCGA COAD/READ were integrated with MSIsensor-based microsatellite instability classification and DepMap CRISPR dependency data from colorectal cancer cell-line models. The analysis focused on the MSS/MSI-L subgroup and aimed to distinguish immune-cold, inflamed, barrier-high and intermediate transcriptomic states, followed by prioritization of candidate tumour-cell-intrinsic and microenvironment-associated vulnerabilities.

### Patient transcriptomic and clinical data source

Colorectal cancer transcriptomic and clinical data were accessed from the publicly available TCGA COAD/READ PanCancer Atlas cohort through cBioPortal for Cancer Genomics [21–23]. The cBioPortal study identifier, molecular profile names, MSIsensor attribute and exported files used for the analysis should be retained with the manuscript records to ensure reproducibility. The study identifier used was: coadread_tcga_pan_can_atlas_2018 RNA-seq expression data were obtained from the corresponding TCGA COAD/READ PanCancer Atlas mRNA expression profile: coadread_tcga_pan_can_atlas_2018_rna_seq_v2_mrn

Gene-expression values were used consistently within the exported cBioPortal profile for within-cohort comparisons. Module scores were derived from z-scored expression values, whereas representative gene-level tables report the expression scale used in the exported analysis files. These values should be interpreted as relative transcriptomic evidence and not as absolute protein abundance or clinical assay positivity.

Sample-level MSIsensor scores were retrieved from cBioPortal clinical annotations using the clinical attribute: MSI_SENSOR_SCORE

Tumours with MSIsensor score <3.5 were classified as MSS/MSI-L and included in the main analysis, consistent with the commonly used MSIsensor threshold for distinguishing microsatellite-stable from microsatellite-instable tumours [24]. Tumours with MSIsensor score ≥3.5 were classified as MSI-H and excluded from this MSS-focused study. Only tumours with both RNA-seq expression data and numeric MSIsensor scores were retained, resulting in 494 MSS/MSI-L colorectal tumours.

### Candidate gene selection

A focused candidate gene set was constructed to represent immune activation, immune suppression, stromal-barrier biology, tumour-intrinsic immune-evasion pathways and druggable cancer vulnerabilities. The selected genes covered the following biological categories: cytotoxic T/NK activity, IFNγ-associated chemokine signalling, antigen presentation, checkpoint/adaptive immune engagement, Treg/myeloid suppression, CAF/ECM barrier biology, WNT/stem-like exclusion, STING/type-I IFN signalling, VEGF/hypoxia/myeloid biology, DNA-damage response, RTK/MAPK/PI3K signalling, epigenetic regulation, apoptosis regulation and clinically actionable colorectal cancer pathways.

The queried candidate set contained 186 genes, of which 183 genes were retrieved from cBioPortal and included in the expression matrix. Three genes, DDX58, MB21D1 and TMEM173, were not returned by cBioPortal under the queried identifiers and were therefore excluded from module scoring where unavailable. Module scores were calculated using the available genes in each module.

### Gene-expression preprocessing and module scoring

Expression values were arranged as a gene-by-sample matrix. For module scoring, expression values for each gene were standardized across samples using z-score transformation. For each sample, module scores were calculated as the mean z-scored expression of the available genes in the corresponding module.

The main modules used for MSS/MSI-L state classification and biological interpretation were:

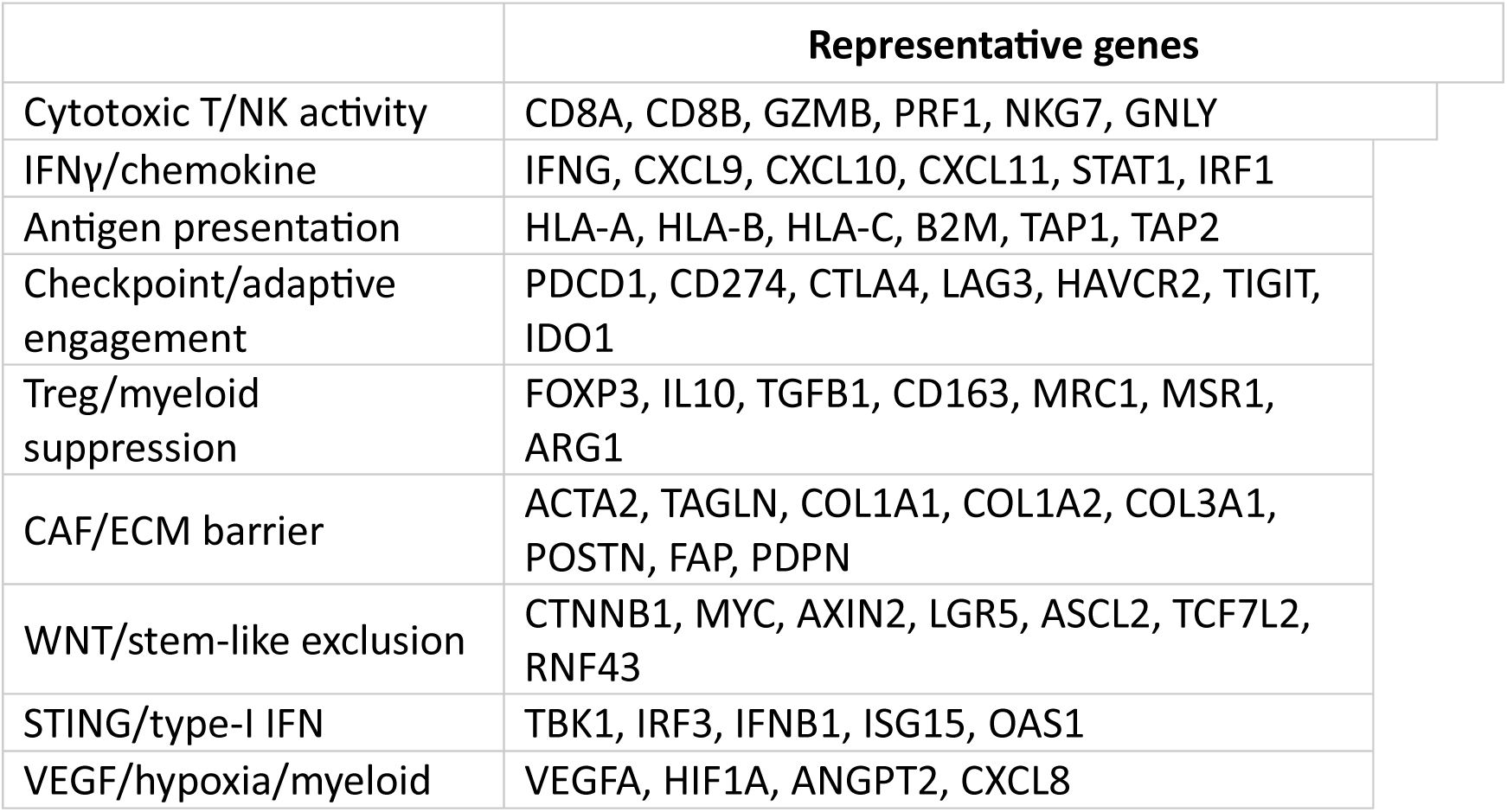

|  | Representative genes |
| --- | --- |
| Cytotoxic T/NK activity | CD8A, CD8B, GZMB, PRF1, NKG7, GNLY |
| IFNγ/chemokine | IFNG, CXCL9, CXCL10, CXCL11, STAT1, IRF1 |
| Antigen presentation | HLA-A, HLA-B, HLA-C, B2M, TAP1, TAP2 |
| Checkpoint/adaptive engagement | PDCD1, CD274, CTLA4, LAG3, HAVCR2, TIGIT, IDO1 |
| Treg/myeloid suppression | FOXP3, IL10, TGFB1, CD163, MRC1, MSR1, ARG1 |
| CAF/ECM barrier | ACTA2, TAGLN, COL1A1, COL1A2, COL3A1, POSTN, FAP, PDPN |
| WNT/stem-like exclusion | CTNNB1, MYC, AXIN2, LGR5, ASCL2, TCF7L2, RNF43 |
| STING/type-I IFN | TBK1, IRF3, IFNB1, ISG15, OAS1 |
| VEGF/hypoxia/myeloid | VEGFA, HIF1A, ANGPT2, CXCL8 |

Additional candidate genes related to DNA-damage response, RTK/MAPK/PI3K signalling, epigenetic regulation and apoptosis regulation were included for target prioritization.

Classification of MSS/MSI-L transcriptomic states

MSS/MSI-L tumours were classified into four relative transcriptomic states using cytotoxic T/NK, IFNγ/chemokine and CAF/ECM barrier module scores:

1. MSS immune-cold
2. MSS hot/inflamed
3. MSS barrier-high
4. MSS intermediate

MSS hot/inflamed tumours were defined by high cytotoxic T/NK and IFNγ/chemokine activity. MSS immune-cold tumours were defined by low cytotoxic T/NK and low IFNγ/chemokine activity. MSS barrier-high tumours were defined by elevated CAF/ECM barrier activity with less dominant cytotoxic immune activation. Tumours not meeting these criteria were assigned to the MSS intermediate group. The classification was cohort-relative and applied to the exported expression and module-score tables. The final sample-level state assignments are provided in the supplementary state table.

This classification was intended as a cohort-relative transcriptomic stratification, not as a clinically validated diagnostic classifier. Consequently, module-level comparisons involving the same scores used for classification were interpreted as descriptive internal checks, whereas candidate prioritization relied additionally on focused gene-level contrasts, DepMap dependency evidence and biological/druggability annotation.

Differential module and gene-level analyses Two main comparisons were performed:

MSS immune-cold vs MSS hot/inflamed MSS barrier-high vs MSS immune-cold

For comparisons where statistical significance or FDR values are reported, module scores and candidate-gene expression values were compared between groups using the Mann-Whitney U test. This non-parametric test was selected because normal distribution of expression values and module scores was not assumed. For each comparison, group means, mean differences, U statistics, nominal P-values and sample sizes were recorded.

### DepMap CRISPR dependency analysis

Functional dependency data were obtained from the Cancer Dependency Map Project using publicly available CRISPR gene-effect data. The analysis used the DepMap Public 26Q1 CRISPRGeneEffect.csv table, which provides post-Chronos gene-effect estimates for all models, with rows corresponding to ModelID and columns corresponding to genes [9,16,20]. The 1208-model matrix used in this study matches the model count listed for DepMap Public 26Q1 gene-dependency summaries. The dependency matrix used was: CRISPRGeneEffect.csv

Model annotations were used to identify colorectal cancer cell-line models. The final dependency analysis included 1208 total cancer models, including 63 colorectal cancer models. For each candidate gene, the mean and median CRISPR gene-effect scores were calculated across colorectal cancer models. More negative gene-effect values indicate stronger growth impairment after gene knockout; a value near 0 indicates little average effect under the screened cell-line conditions. A CRC dependency score was defined as the negative of the mean gene-effect value in colorectal cancer models, so that higher positive values indicate stronger average dependency.

The following dependency metrics were summarized for each candidate gene: number of colorectal cancer models, mean CRC gene-effect score, median CRC gene-effect score, proportion of CRC models with gene effect <−0.5, proportion of CRC models with gene effect <−1.0, and dependency behaviour across all cancer models. Genes showing broad dependency across many cancer types were interpreted cautiously as common essential genes rather than CRC-specific vulnerabilities.

### Candidate target prioritization

Candidate targets were prioritized by integrating four evidence layers in a descriptive, hypothesis-generating framework:

1. patient transcriptomic evidence for immune-cold association,
2. patient transcriptomic evidence for barrier-high association,
3. DepMap colorectal cancer dependency evidence, and
4. biological/druggability annotation.

Cold-associated expression evidence was derived from the MSS immune-cold versus MSS hot/inflamed comparison. Barrier-associated expression evidence was derived from the MSS barrier-high versus MSS immune-cold comparison. DepMap evidence was derived from the CRC dependency score. Candidate genes were also annotated for immune relevance and druggability based on known biological function and therapeutic relevance in cancer or colorectal cancer.

Druggability categories included clinically actionable targets, druggable pathways, epigenetic regulators, mutation-specific emerging targets and genes with uncertain druggability. This annotation was used for prioritization only and was not interpreted as evidence of clinical efficacy. Integrated scores were not trained against patient outcome, survival or immunotherapy-response endpoints. The integrated ranking was used as a hypothesis-generating framework to distinguish tumour-cell-intrinsic vulnerabilities from patient-derived microenvironmental barrier programmes.

### Data visualization

Figures were prepared to summarize MSS/MSI-L state distribution, immune and stromal module scores, focused gene-level differences, DepMap dependency evidence and integrated target prioritization. Descriptive panels show distributions or rankings; statistical significance is shown only where FDR-adjusted testing was performed.

A schematic model was also generated to summarize the proposed state-matched therapeutic framework.

### Statistical approach

For analyses where significance is reported, two-group comparisons used the Mann-Whitney U test and Benjamini-Hochberg correction for multiple testing. DepMap dependency summaries and integrated prioritization scores were descriptive and hypothesis-generating.

## Ethics statement

This study used publicly available, de-identified data from TCGA/cBioPortal and DepMap. No new human samples were collected, and no identifiable patient information was used. Therefore, additional institutional ethics approval was not required.

## Author contributions

A.T. conceptualized the manuscript and performed data analysis. D.N. retrieved public datasets and performed analysis and preparation of figures. All authors were involved with writing and revision of manuscript.

## Data availability statement

The main processed output tables and generated figures are provided as supplementary files. TCGA COAD/READ expression and clinical data are publicly available through cBioPortal under the study identifier: coadread_tcga_pan_can_atlas_2018 DepMap CRISPR gene-effect and model annotation files are publicly available through the DepMap portal. The analysis used: CRISPRGeneEffect.csv, Model.csv or sample_info.csv

---

**Supplementary Figure 1.**
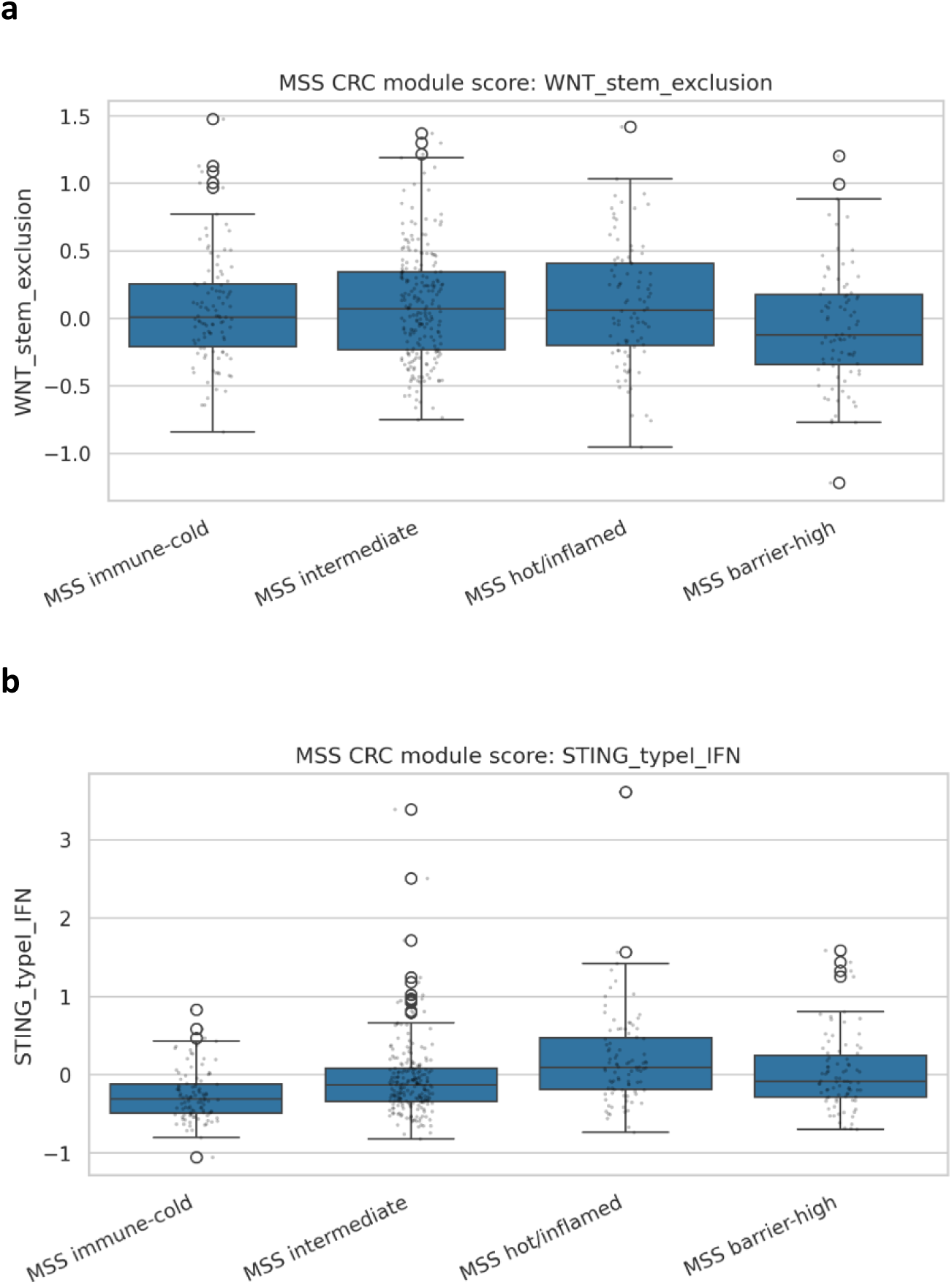
WNT/stem-like exclusion and STING/type-I IFN module scores across MSS/MSI-L transcriptomic states: Additional curated module scores were compared across MSS immune-cold, MSS hot/inflamed, MSS barrier-high and MSS intermediate tumours. WNT/stem-like exclusion and STING/type-I IFN module scores were calculated as the mean z-scored expression of available module genes. These exploratory modules were included to assess tumour-intrinsic exclusion and innate immune/type-I interferon-associated programmes across MSS states.

**Supplementary Figure 2. Expanded integrated target ranking.** Expanded visualization and tabular ranking of the top integrated candidate vulnerabilities are provided as supplementary files. Expanded visualization of the top candidate genes from the integrated prioritization analysis. Candidate ranking incorporated patient transcriptomic evidence for immune-cold or barrier-high association, DepMap colorectal cancer dependency evidence, immune relevance, druggability annotation and common-essentiality penalty. Genes with strong stromal patient-signature evidence but limited tumour-cell-line dependency were interpreted as microenvironmental markers rather than tumour-cell-intrinsic vulnerabilities.

